# Holistic Characterization of Single Hepatocyte Transcriptome Responses to High Fat Diet

**DOI:** 10.1101/2020.04.16.045260

**Authors:** Sung Rye Park, Chun-Seok Cho, Jingyue Xi, Hyun Min Kang, Jun Hee Lee

**Affiliations:** Department of Molecular and Integrative Physiology and Institute for Gerontology, University of Michigan Medical School, Ann Arbor, MI 48109, USA; Department of Biostatistics and Center for Statistical Genetics, University of Michigan School of Public Health, Ann Arbor, MI 48109, USA

**Keywords:** scRNA-seq, hepatocytes, obesity, NAFLD, HFD

## Abstract

During nutritional overload and obesity, hepatocyte function is grossly altered, and a subset of hepatocytes begins to accumulate fat droplets, leading to non-alcoholic fatty liver disease (NAFLD). Recent single cell studies revealed how non-parenchymal cells, such as macrophages, hepatic stellate cells, and endothelial cells, heterogeneously respond to NAFLD. However, it remains to be characterized how hepatocytes, the major constituents of the liver, respond to nutritional overload in NAFLD. Here, using droplet-based single cell RNA-sequencing (Drop-seq), we characterized how the transcriptomic landscape of individual hepatocytes is altered in response to high-fat diet (HFD) and NAFLD. We showed that the entire hepatocytes population undergoes substantial transcriptome changes upon HFD, although the patterns of alteration were highly heterogeneous with zonation-dependent and –independent effects. Periportal (zone 1) hepatocytes downregulated many zone 1-specific marker genes, while a small number of genes mediating gluconeogenesis were upregulated. Pericentral (zone 3) hepatocytes also downregulated many zone 3-specific genes; however, they upregulated several genes that promote HFD-induced fat droplet formation, consistent with findings that zone 3 hepatocytes accumulate more lipid droplets. Zone 3 hepatocytes also upregulated ketogenic pathways as an adaptive mechanism to HFD. Interestingly, many of the top HFD-induced genes, which encode proteins regulating lipid metabolism, were strongly co-expressed with each other in a subset of hepatocytes, producing a variegated pattern of spatial co-localization that is independent of metabolic zonation. In conclusion, our dataset provides a useful resource for understanding hepatocellular alteration during NAFLD at single cell level.

## Introduction

The liver is a vital organ that performs essential digestive and metabolic functions within the body, such as glucose and fat metabolism, serum protein production, bile secretion, and chemical detoxification. Most of these functions are mediated by hepatocytes, which constitute the major cell type of the liver, and comprise of 70-85% of the liver’s mass.

With the prevalence of obesity in the modern society, the incidence of non-alcoholic fatty liver disease (NAFLD) is increasing at an alarming rate (25). During obesity, over-nutrition and sedentary lifestyle lead to a chronic calorie surplus, resulting in the storage of excessive nutrients in the form of fat. In this condition, the liver also accumulates large fat droplets, which does not typically occur in healthy liver. NAFLD can precipitate further advanced liver diseases such as steatohepatitis (NASH), liver cirrhosis, and hepatocellular carcinoma (HCC, liver cancer) (30).

Recently, pathological NAFLD responses of non-parenchymal cell types, such as inflammatory cells and hepatic stellate cells, were characterized at single cell level through scRNA-seq (52). It was reported that, during NAFLD, some macrophages and hepatic stellate cells still retain their normal transcriptome that is almost indistinguishable from those in healthy liver. However, new cell types, which have activated inflammatory signaling (NASH-associated macrophages) or fibrogenic responses (activated hepatic stellate cells), emerged from the normal population and occupied a substantial portion of cells in the diseased liver. Similar observations were made from fibrotic responses of hepatic stellate cells to carbon tetrachloride treatment (7, 24) or human liver cirrhosis (34), indicating the presence of both resting and activated hepatic stellate cell population in fibrotic liver. Another recent study indicated that liver endothelial cells also show similar bipartite response to NASH with responsive and unresponsive populations (16). These findings suggested that at least some non-parenchymal liver cells maintain unaltered transcriptome phenotypes to mediate homeostatic function, while other cells alter their transcriptome or migrate from other places to play either adaptive or maladaptive pathological roles during NASH or NAFLD.

Although hepatocytes are often considered functionally homogeneous, studies actually indicate that individual hepatocytes are exposed to different physiological environments, receive different developmental cues, express different sets of genes, and thereby play specialized metabolic functions according to their histological niche (4, 18-20). Recent single cell RNA-seq (scRNA-seq) studies on normal mouse and human liver samples confirmed the presence of such heterogeneity in mammalian liver (1, 28). Furthermore, histological studies revealed that a subset of hepatocytes in a specific region is more prone to fat accumulation (NAFLD) (5, 14, 15), fibrotic disease progression (NASH) (10), liver damage, and hepatocarcinogenesis (HCC) (40, 43, 48). Therefore, although transcriptomic analyses of bulk liver mRNAs have revealed that lipogenic, glucogenic, and inflammatory gene transcription levels substantially change upon the development of NAFLD and NASH (3), it is unknown how individual hepatocytes alter their gene expression during liver pathogenesis.

Here we performed droplet-based single cell RNA-sequencing (Drop-seq) (27) on hepatocytes freshly isolated from lean and high fat diet (HFD)-fed obese mice and characterized their single cell transcriptome. Our analyses indicate that, unlike non-parenchymal cell types that have both non-responsive and responsive populations, all hepatocytes altered their transcriptome upon HFD, and each of their single cell transcriptome was distinct from the ones isolated from lean mice. However, the patterns of transcriptome alteration were highly heterogeneous across the metabolic zones, and there are also HFD response heterogeneity that is independent of the zonation profile. Some of these interesting single cell gene expression features were observed at the protein level through immunohistochemistry of liver sections. Collectively, our work reveals how HFD alters the transcriptomic landscape of single hepatocytes across the whole population.

## Materials and Methods

### Data Availability

The scRNA-seq dataset generated from this study is available at the Gene Expression Omnibus database (GEO Accession number: GSE157281). The data can also be accessed through an interactive online resource (https://lee.lab.medicine.umich.edu/hfd), which has an intuitive graphical user interface for exploring our scRNA-seq dataset.

### Mice and Diets

8-week-old C57BL/6J littermate male mice were separated into two groups and were fed on a regular chow diet (LFD group; Lab Diet, 5L0D) or high fat diet (HFD group; Bio-Serv, S3282). After 12 weeks of dietary modulation, mice whose body weight reached between 48 and 52 g (HFD group) or between 35 and 38 g (LFD group) were euthanized and subjected to single hepatocyte isolation and Drop-seq. We complied with all relevant ethical regulations for animal testing and research. All experiments were approved by the University of Michigan Institutional Animal Care & Use Committee (PRO00007710 and PRO00009630).

### Hepatocyte isolation

For hepatocyte isolations, the liver was first perfused with calcium-free Hank’s Balanced Salt Solution (HBSS; 14175-095, Gibco) containing 0.2 mg/mL EDTA (51201, AccuGENE) and sodium bicarbonates (7.5%; 25080-094, Gibco) and then sequentially perfused with 0.2% collagenase type II (LS004196, Worthington) in Hank’s Balanced Salt Solution (HBSS; 14025-092, Gibco) with Calcium Chloride (2.5M; C7902-500G, Sigma). The collagenase-treated liver was extracted from the body and further incubated at 37°C for 20 min. Liver cells were diluted in DMEM (11965, Gibco) containing 10% serum and centrifuged at 50 g for 5 min to enrich hepatocytes and passed through a 100-micron nylon cell strainer (10199-659, VWR) multiple times. To remove non-hepatocytes, the gradient precipitation using a 30% percoll solution (17-5445-02, GE Healthcare) was performed, and the resulting hepatocytes were resuspended in 0.5% BSA (A8806, Sigma) in PBS (11965-092, Gibco) for the further analysis of viability and subsequent Drop-Seq experiment.

### Drop-seq library preparation

Drop-seq experiments were performed through a previously described method (27). Hepatocyte preparations were diluted in 2 ml PBS-BSA to a final concentration of 240,000∼300,000 cells. Diluted hepatocytes suspension, barcoded beads (MACOSKO-2011-10, Chemgenes) in lysis buffer (400 mM Tris pH7.5, 40 mM EDTA, 12% Ficoll PM-400, 0.4% Sarkosyl and 100mM DTT; 100,000 beads/ml), and droplet generation oil (184006, Bio-rad) were injected into a microfluidics device (FJISUM-QO-180221, FlowJEM) through three separate inlets. The flow rates for the cell and bead suspensions were set as 2,000 µl/hr, and the flow rate for the droplet oil was set to 7,500 µl/hr. Resulting droplets were sequentially collected in 50 ml falcon tubes, and the total collection time was between 25 and 30 mins. Droplets were broken by vigorous shaking to release the beads into the solution, and the beads were collected by centrifugation. Beads were washed multiple times in 6X SSC (diluted from 20X SSC; 15557044, Invitrogen). Excess bead primers were removed by the treatment of Exonuclease I (NEBM0293S, NEB), cDNA synthesis was performed using Template Switch Oligo (TSO), and DNA was amplified using PCR, according to the original Drop-seq protocol (27). The resultant PCR product was purified by AMPure XP beads (A63881, Beckman Coulter). The products of the multiple PCR reactions were used for the secondary PCR to construct a full-length cDNA library, which was processed into the sequencing library using the Nextera XT DNA Library Preparation Kit (FC-131-1096, Illumina) with unique barcode sequences for each set. The quality of the libraries was inspected by agarose gel electrophoresis for their average size and concentration before pooling for the sequencing. A total of 5 sets of cDNA libraries from Drop-seq runs, two from LFD liver, and three from HFD liver, were analyzed. The pooled libraries were sequenced using Illumina HiSeq-4000 High-Output at the UM Sequencing Core and AdmeraHealth Inc. after an additional quality control process through Agilent BioAnalyzer.

### Drop-seq data processing

We processed raw reads following the instructions described in the Drop-seq Laboratory Protocol v3 (27) using DropSeqTools (v1.13). Reads were aligned to the mm10 mouse genome using STAR (v.2.6.0a) (8) following the default DropSeqTools pipeline. The aligned reads were further processed using a *popscle* software tool (https://github.com/statgen/popscle) to produce the digital expression matrix. We used a unique molecular identifier (UMI) count 400 as an initial cutoff to filter 44,245 droplets to consider for more stringent filtering. Because hepatocytes are extremely fragile (38, 42), ambient RNAs (soup) released from dead hepatocytes could be easily captured by the majority of droplets that do not have actual single cells. Indeed, preliminary analysis of Drop-seq results revealed several droplet clusters that were suspected of having been formed from soup, not from a single cell. To identify these soup droplets from our dataset, a shuffled (Shf) dataset was generated by random shuffling of transcriptome information in the original (Org) dataset. We assumed that soup droplets in the Org dataset would exhibit characteristics similar to the droplets in the Shf dataset. To test this, Org and Shf dataset were plotted on the t-SNE manifold. Indeed, the results indicate that many of the Org droplets from the Drop-seq experiments have a characteristic similar to droplets of the Shf dataset, as they overlap in the t-SNE manifold (Fig. S1A). From the t-SNE manifold, we identified four small clusters (Fig. S1A) that are unique to the Org dataset. Among these, one cluster (cluster AA in Fig. S1A and S1B) contained higher levels of mitochondrial transcripts while another cluster (cluster BB in Fig. S1A and S1C) contained very low levels of UMI. The other two clusters (clusters CC and DD in Fig. S1A-S1D) had relatively higher UMI numbers and relatively lower mitochondrial transcript content; therefore, we focused on isolating these two clusters from the dataset. Therefore, through a series of multi-dimensional clustering and subtraction of irrelevant droplets with soup-like profiles (Shf-enriched clusters), higher mitochondrial contents (cutoff: 30%) and lower UMI counts (cutoff: 1000), we isolated a total of 454 droplets that represent 216 cells from two LFD liver samples and 238 cells from three HFD liver samples (Fig. S1D).

### Cell clustering and data visualization

The digital expression matrix was processed to Seurat v3 (41) following the “standard processing workflow” in the tutorial. 2-dimensional t-SNE (45) and UMAP (2) manifolds were used to visualize gene expression data across different clusters of single cells. Clustering was performed using the shared nearest neighbor modularity optimization implemented in Seurat’s *FindClusters* function using a resolution parameter as 0.2. We observed that batch effects are minimal, and all HFD droplets across three independent batches fell into the cluster corresponding to HFD cells, while most of the LFD droplets (97%) across two independent batches fell into the other cluster corresponding to LFD cells.

### Imputation of single cell expression

We performed the imputation of the data using *magic* package (47) or using *saver* package (17). Default parameters were used for the imputation work. The scatterplots and feature plots of imputed data were visualized using customized R scripts with *ggplot2*.

### Construction of hepatocyte zonation profiles

*Arg1* and *Cyp2e1* are established markers for zone 1 and 3 hepatocytes, respectively (1, 12), and expression levels of these genes were comparable between LFD and HFD livers in our dataset. Accordingly, imputed gene expression levels for *Arg1* and *Cyp2e1* were used for estimating hepatocyte zonation. Zonation score was calculated as the difference between the *magic*-imputed levels of *Arg1* and *Cyp2e1* expression. According to the zonation score, hepatocytes were divided into five bins of cells, among which the top 3 bins were grouped together as zone 1 hepatocytes, and the bottom two bins were grouped as zone 2 and 3 hepatocytes, respectively. The resultant hepatocyte groups appropriately reflect the biological characteristics of zone 1-3 hepatocytes, as supported through independent visualizations using PCA, t-SNE and UMAP manifolds, as well as gene expression analyses of the other established zone-specific markers (see Results and Discussion for details).

### Pathway enrichment analysis

Differentially expressed genes (based on fold-enrichment) were identified between LFD and HFD hepatocytes, and between Zone 1 and Zone 3 hepatocytes from the LFD set of hepatocytes, using *FindAllMarkers* function in Seurat. Networks of GO terms were constructed using ShinyGO v0.61 (11), using only the top 20 significant terms. The pathway enrichment analysis was also performed using *enrichGO* and *enrichKEGG* functions in the clusterProfiler version 3.6 (53).

### Immunohistochemistry

For immunohistochemistry, liver tissues were fixed in 10% buffered formalin and embedded in paraffin and subjected to immunohistochemical staining, as previously described (6). In brief, paraffin-embedded liver sections were incubated with primary antibodies obtained from Santa Cruz Biotechnology (Apoa4, sc-374543; Elovl5, sc-374138; Fabp1, sc-271591; Cyp2f2, sc-374540; Cyp1a2, sc-53241) at 1:100, followed by incubation with biotin-conjugated secondary antibodies (Vector Lab, BA-9200; 1:200) and horseradish peroxidase (HRP)-conjugated streptavidin (BD Biosciences, 554066; 1:300). The HRP activity was visualized with diaminobenzidine staining, and nuclei were visualized by Haematoxylin counterstaining. For fluorescence staining of lipid droplets and Cyp2f2, livers were harvested from 4 month-old mice, which had been either LFD or HFD for two months. Frozen liver sections were fixed with 2% paraformaldehyde, blocked with 1X Western Blocking Reagent (Roche), and incubated with anti-Cyp2f2 primary antibody (sc-374540; Santa Cruz Biotechnology), followed by Alexa 594-conjugated secondary antibody, DAPI and BODIPY 493/503 (Invitrogen).

## Results and Discussion

### Drop-seq successfully captures single hepatocyte transcriptome profile

To characterize the effect of HFD on single cell transcriptome of hepatocytes, we performed Drop-seq in five independent experiments with freshly isolated hepatocytes from two normal chow (low fat diet; LFD)-fed lean mice and three HFD-fed obese mice. A total of 216 high-quality hepatocytes were identified from the livers of LFD mice, while 238 were identified from those of HFD mice (see Materials and Methods for details). All of these droplets expressed robust levels of *Alb* (>0.9% of total transcriptome; Fig. S2A), an authentic hepatocyte marker encoding albumin protein, confirming that these droplets indeed represent hepatocyte transcriptome.

In contrast, most macrophage markers, such as *Emr1, Itgam* and *Cd14*, as well as many inflammatory cytokines, such as *Tnf, Il6* and *Ccl2*, were undetectable from our single cell transcriptome dataset (Fig. S2B and S2C), indicating that our Drop-seq preparations did not have contaminating fractions of Kupfer cells, the liver-resident macrophages. Many markers for hepatic stellate cells and fibroblasts, such as *Acta2, Col3a1, Pecam1*, and *Mmp2*, were also not detected (Fig. S2D). Major adipocyte markers, such as *Fabp4* and *Adipoq*, were also undetectable (Fig. S2E), indicating that, although HFD and fatty liver can render hepatocytes to accumulate lipid droplets (32), they do not alter the tissue identity of hepatocytes to exhibit adipocyte characteristics.

### HFD alters single cell transcriptome profile of entire hepatocyte population in liver

To explore and understand the single hepatocyte transcriptome data, we first performed the principal component (PC) analysis to determine the signatures of the largest variance in our dataset. PC1, which represents the largest variance, did not characterize significant differences between LFD and HFD samples (Fig. 1A, left). However, PC2 and PC3, the orthogonal axes representing the second and third largest variance, respectively, were highly effective in separating LFD and HFD transcriptome profiles (Fig. 1A, center and right). Correspondingly, PC2 and PC3 were sufficient to discriminate LFD and HFD hepatocytes without any additional information (Fig. 1B). The effect of HFD was robust across independent batches of the experiment (Fig. 1C).

**Fig. 1.**
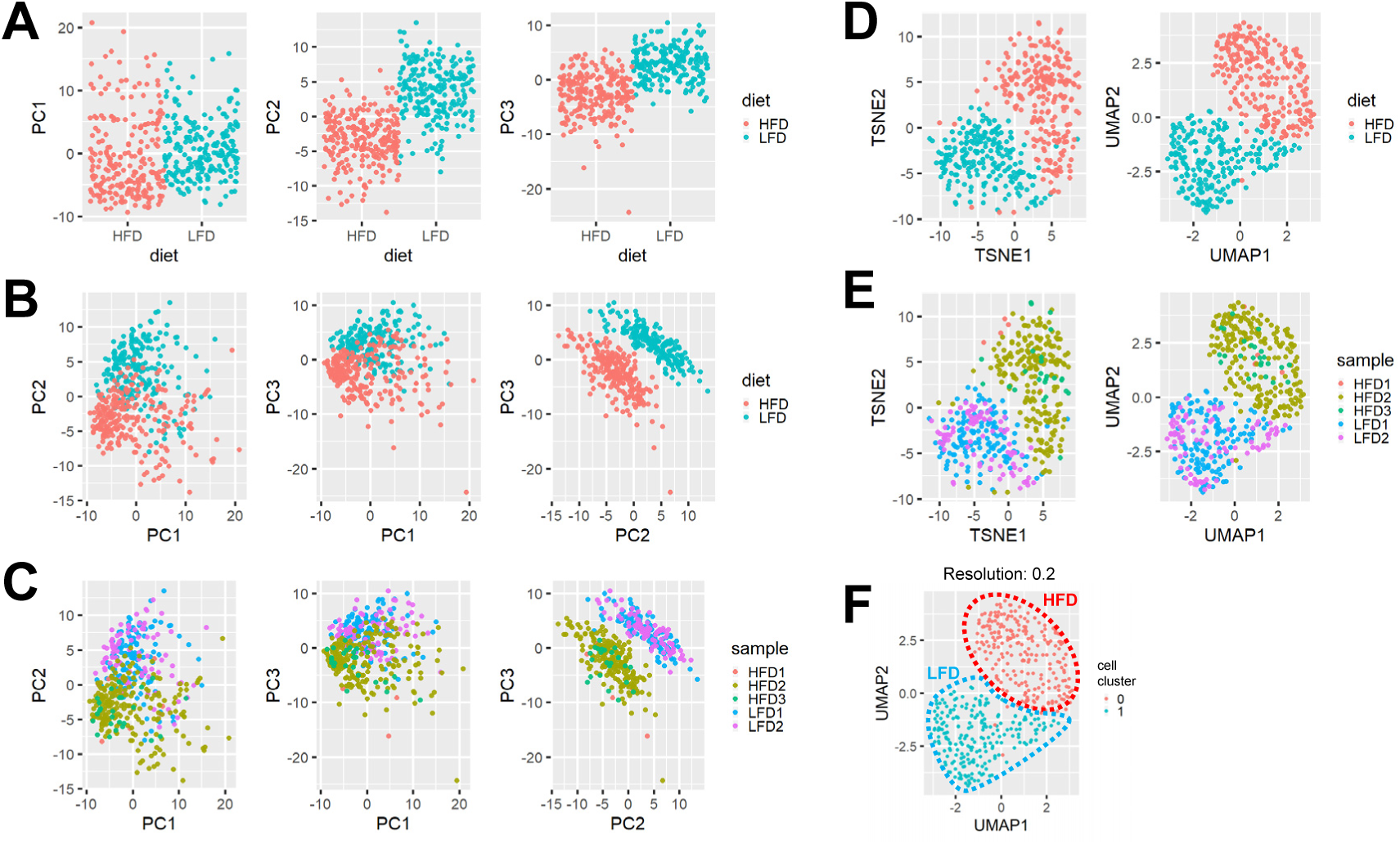
HFD alters single cell transcriptome of the entire hepatocyte population. 8-week-old C57BL/6J male littermate mice were separated into two groups and were fed on a regular chow diet (LFD group) or high fat diet (HFD group). Drop-seq of hepatocytes was performed after 12 weeks of dietary modulation. (A-F) Principal component analysis (PCA; A-C), t-SNE, and UMAP (D-F) manifolds, colored with diet (A, B and D), sample (C and E), or the result from multi-dimensional clustering (F). Individual dots represent single cell transcriptome. In each manifold, the distance between individual dots represents the difference between the single cell transcriptome. Approximate boundaries of the area for LFD and HFD samples were indicated as dotted outline (F).

Similar trends were observed from the nonlinear manifolds generated by t-SNE and UMAP dimension reduction methods (2, 46), which segregated LFD and HFD groups of hepatocytes (Fig. 1D) but robust against batch effects (Fig. 1E). High-dimensional clustering analysis also clearly differentiated the LFD and HFD groups; all cells from HFD mice fell into the cluster corresponding to HFD group (group 0 in Fig. 1F), while 213 out of 216 cells (98%) from LFD mice fell into the LFD group (group 1 in Fig. 1F). These results indicate that the entire hepatocyte population in mouse liver responded to the HFD challenge by altering their transcriptome profiles.

### Metabolic zonation of hepatocytes was captured in both HFD and LFD livers

We were curious about the nature of the PC1 axis, which represents the largest variance of transcriptomic profiles across all hepatocyte populations, yet does not strongly represent the diet effect (Fig. 1A). We observed that the hepatocyte marker *Alb* expression exhibited a substantial negative correlation with PC1 in both HFD and LFD groups (Fig. S2F; *r* = -0.54, *P <* 2.2e-16). *Alb* expression is known to be relatively higher in periportal zone 1 hepatocytes and relatively lower in pericentral zone 3 hepatocytes (9); therefore, we suspected that the PC1 axis might represent the metabolic zonation of individual hepatocytes. To further substantiate this conjecture, we examined the expressions of well-characterized periportal marker *Arg1* and pericentral marker *Cyp2e1* (1, 12) to understand the zonation structure of our dataset. Both scaled and imputed expression levels of *Arg1* showed negative correlation with the PC1 (Fig. S3A, upper; *r* = -0.26 and -0.84, respectively; *P* < 2.5e-8) while expression levels of *Cyp2e1* showed positive correlation with the PC1 (Fig. S3A, lower; *r* = 0.56 and 0.89; *P* < 2.2e-16), indicating that PC1 indeed represents the metabolic zonation structure of hepatocytes.

Interestingly, in single cells, imputed levels of *Arg1* and *Cyp2e1* expression showed a clear negative correlation (Fig. 2A; *r* = -0.93), consistent with their opposed expression patterns in the liver (1, 12). Based on these levels of *Arg1* and *Cyp2e1* expression, we partitioned the liver with three zones: zone 1 with periportal characteristics, zone 2 with intermediate characteristics, and zone 3 with pericentral characteristics (Fig. 2A, right).

**Fig. 2.**
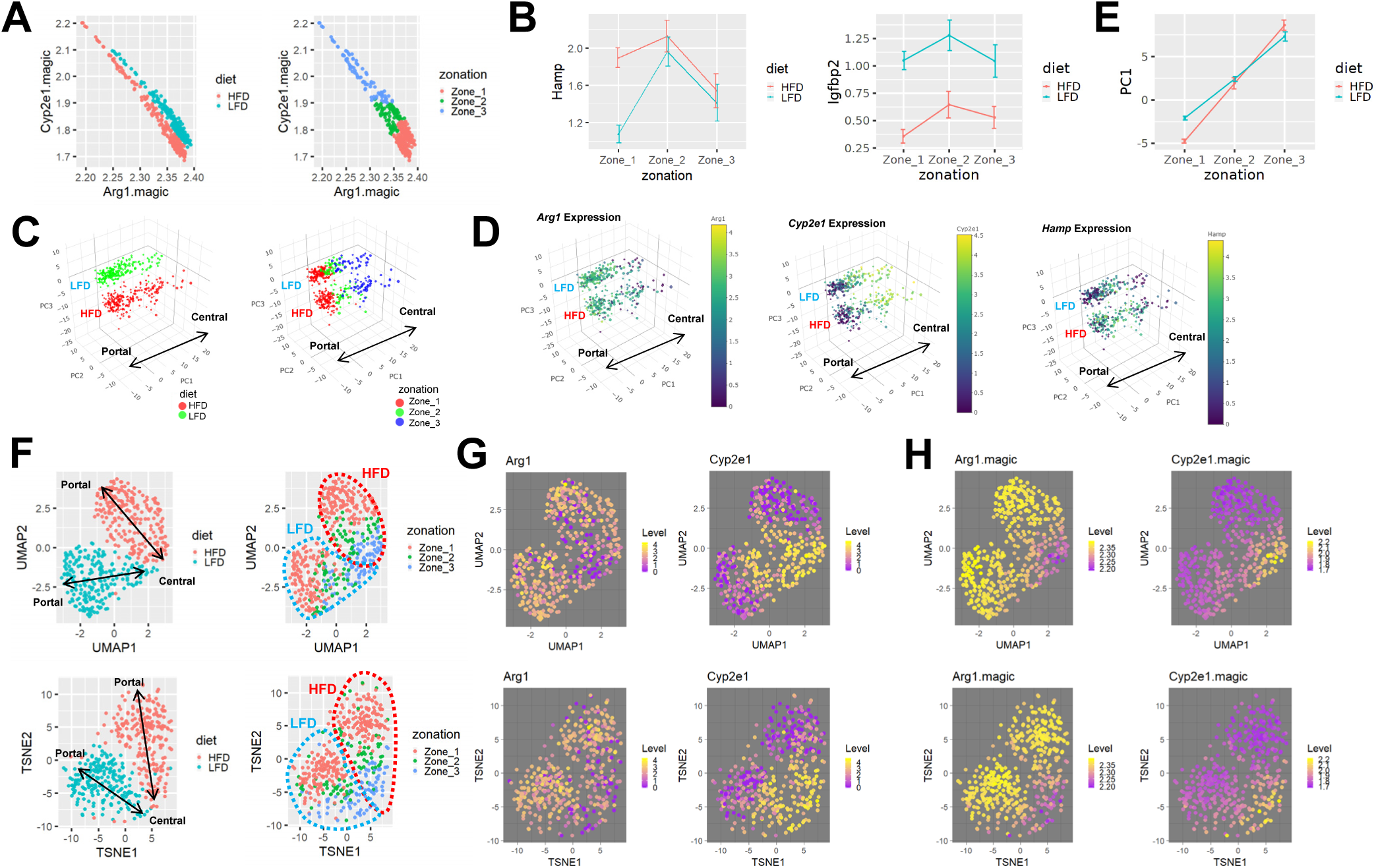
Zonation patterns of single hepatocyte transcriptome is preserved after HFD. (A) Inverse correlation between imputed expression levels of *Arg1* and *Cyp2e1* (*magic*-imputed expression values). Individual dots represent single cell transcriptome, colored with diet (left) and zone assignment (right). (B) Analysis of single cell gene expression in hepatocytes of each zone, expressed as mean±SEM (scaled expression values). Data from LFD and HFD livers were analyzed separately. (C and D) 3-dimensional PCA manifolds depicting the effect of diet (C, left), zonation (C, right), and expression levels of indicated genes (D). Individual dots represent single cell transcriptome. The size of dots represent the number of RNA features captured in the droplet. PC1 is composed of genes showing zone-specific expression patterns. PC2 and PC3 are composed of genes showing diet-regulated expression patterns. LFD and HFD area, as well as the directionality of metabolic zonation (from portal to central), are indicated in each manifold. (E) Analysis of single cell PC1 values in hepatocytes of each zone, expressed as mean±SEM (raw component scores). Data from LFD and HFD livers were analyzed separately. (F-H) UMAP (top) and t-SNE (bottom) manifolds depicting the effect of diet (F, left), zonation (F, right), and scaled (G) and imputed (H) expression levels of indicated genes. LFD and HFD area (F, right), as well as the directionality of metabolic zonation (from portal to central; F, left), are indicated.

Previous studies isolated a long list of zone 1-specific markers, such as *Alb, Ass1, Arg1, Cyp2f2, Cps1, Gls2, Pck1* and *Sult5a1*, and zone 3-specific markers, such as *Glul, Oat, Slc1a2, Lect2, Ldhd, Por, Cyp1a2, Cyp2e1, Ahr*, and *Gstm2, 3* and *6*, through various methodologies including differential isolation, immunohistochemistry or RNA *in situ* hybridization analyses (1, 12, 44). All of these genes appear to have corresponding patterns of expression in our dataset (Fig. S4). Furthermore, *Hamp* and *Igfbp1*, genes whose expression is elevated in the intermediate region of the liver (1), showed zone 2-specific expression from our dataset (Fig. 2B). These results confirm the validity of our zonation method.

Diet and zonation effects can also be jointly visualized in 3-dimensional PC1/PC2/PC3 space, where PC2 and PC3 axes separate LFD and HFD hepatocytes (Fig. 2C, left), and the PC1 axis visualizes the portal-to-central histological zonation structure (Fig. 2C, right; Fig. 2D and S3B). Indeed, PC1 values were the highest in zone 3 and the lowest in zone 1 hepatocytes, according to our hepatocyte zonation groups (Fig. 2E). Diet and zonation effects were also robustly observed in UMAP and t-SNE (Fig. 2F-2H) manifolds. These results indicate that (i) the structure of metabolic zonation is maintained in HFD liver, (ii) HFD produced transcriptome-altering effects on hepatocyte population across entire zonation niches, and (iii) zonation effect and diet effect are the two major sources of variation in single hepatocyte transcriptomes observed from our dataset.

### HFD alters the expression of genes controlling lipid metabolism

Using the diet and zonation information of individual hepatocytes, we identified a list of genes whose expression patterns are modulated by HFD or dependent on their zonation. 91 genes were significantly upregulated in the HFD group, while 226 genes were significantly upregulated in the LFD group (FDR<0.01; Table S1, first to third tabs). Partially overlapping with this list (Fig. 3A), 74 genes were found to be specific to zone 1 hepatocytes, while 320 genes were specific to zone 3 hepatocytes (FDR<0.01; Table S1, first, fourth and fifth tabs). Heat map analysis of the diet-specific (Fig. 3B) and zone-specific genes (Fig. 3C) confirmed that these gene groups show contrasted gene expression patterns across different populations of hepatocytes.

**Fig. 3.**
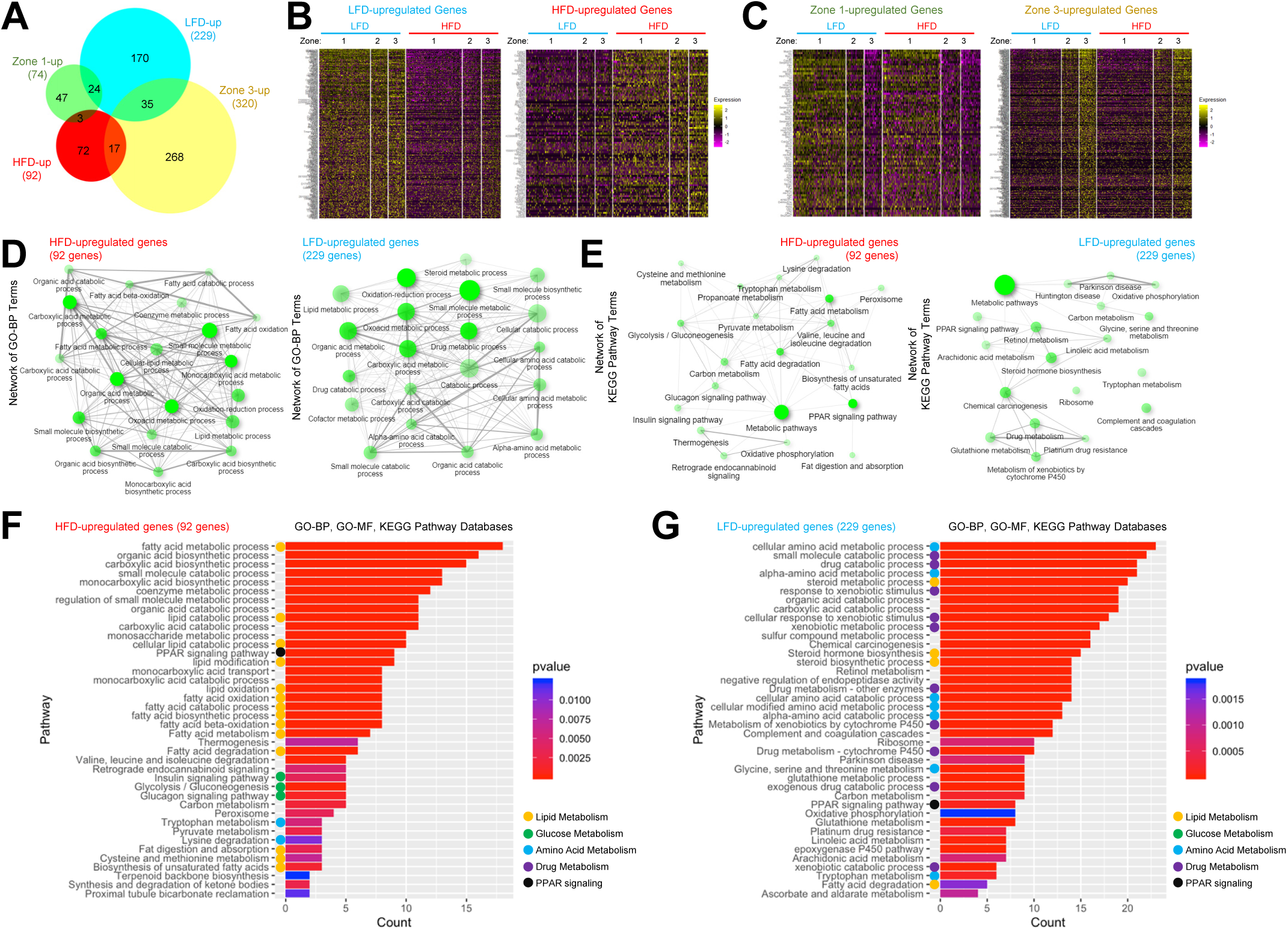
Isolation of genes showing diet- and zone-specific expression patterns. (A) Area-proportional Venn diagram depicting the relationship between diet-regulated genes and zone-specific genes. (B and C) Heat map analysis depicting gene expression across single cell population. Single cells were clustered into six groups (thick columns) according to diet and zone. Diet-regulated genes (B) and zone-specific genes (C) were analyzed. (D and E) Network analysis of gene ontology-biological pathway (GO-BP; D) and Kyoto Encyclopedia of Genes and Genomes (KEGG; E) pathway enrichment terms, using ShinyGO (11). Pathways whose enrichment is significant (FDR < 0.05; top 20 terms) were presented as nodes. Two nodes are connected if they share 20% or more genes. Darker nodes are more significantly enriched gene sets. Bigger nodes represent larger gene sets. Thicker edges represent more overlapped genes. (F and G) Enrichment analysis of HFD-upregulated (left) and downregulated (right) genes, using clusterProfiler (53) with GO-BP, GO-molecular function (GO-MF) and KEGG databases. Color of bars indicates significance (*P* values) while length of bars indicates gene count. Color of circles indicate GO terms related to lipid metabolism (yellow), glucose metabolism (green), amino acid metabolism (blue), drug metabolism (purple) and PPAR pathway (black).

Gene ontology analysis of HFD-upregulated and -downregulated (LFD-upregulated) genes showed that, consistent with previous bulk gene expression studies (22, 37, 39), genes controlling lipid and fatty acid metabolism are upregulated in HFD, while genes controlling amino acid and drug catabolism are downregulated (Fig. 3D-3G; FDR<0.05 for all presented pathways). However, genes mediating inflammation and fibrosis were not included here (Fig. 3D-3G; Table S1), since our dataset was exclusive to hepatocytes and did not include hepatic stellate cells or inflammatory cells (Fig. S1). Pathway enrichment analysis using the Kyoto Encyclopedia of Genes and Genomes (KEGG) database identified the metabolic pathways as the top enriched pathway for both HFD-upregulated and -downregulated gene lists (Fig. 3E; FDR = 1.6e-10 and 2.2e-29, respectively), consistent with the central role of the liver in metabolism. Interestingly, among various biological pathways, the PPAR pathway was represented in both HFD-upregulated and HFD-downregulated gene lists (Fig. 3E-3G), consistent with the former studies indicating that the pathway is among the major pathways altering hepatocellular transcriptome during HFD (33, 49).

### HFD alters expression patterns of a subset of zone-specific genes

Next, we focused on the genes that exhibit both diet- and zone-specific expression patterns (Fig. 3A; Table S1, first tab). We found that many markers for zone 1 hepatocytes, such as *Cyp2f2, Mup1, Gstp1* and *Hpx*, were strongly downregulated upon HFD feeding (Fig. 4A-4D). Only a very small number of zone 1-specific genes, such as *Aldob, Fbp1* and *Mup21*, were upregulated (Fig. 4E and 4F). Many zone 3 hepatocyte markers, such as *Cyp1a2, Mup17, Gstm1* and *Cyp2a5*, were also downregulated (Fig. 4A-4D). However, several zone 3-specific genes, including *Cyp4a14, Aldh3a2*, and *Csad*, were substantially upregulated in response to HFD (Fig. 4E and 4G). Therefore, the HFD effect on zone-specific gene expression could be variable across individual genes.

**Fig. 4.**
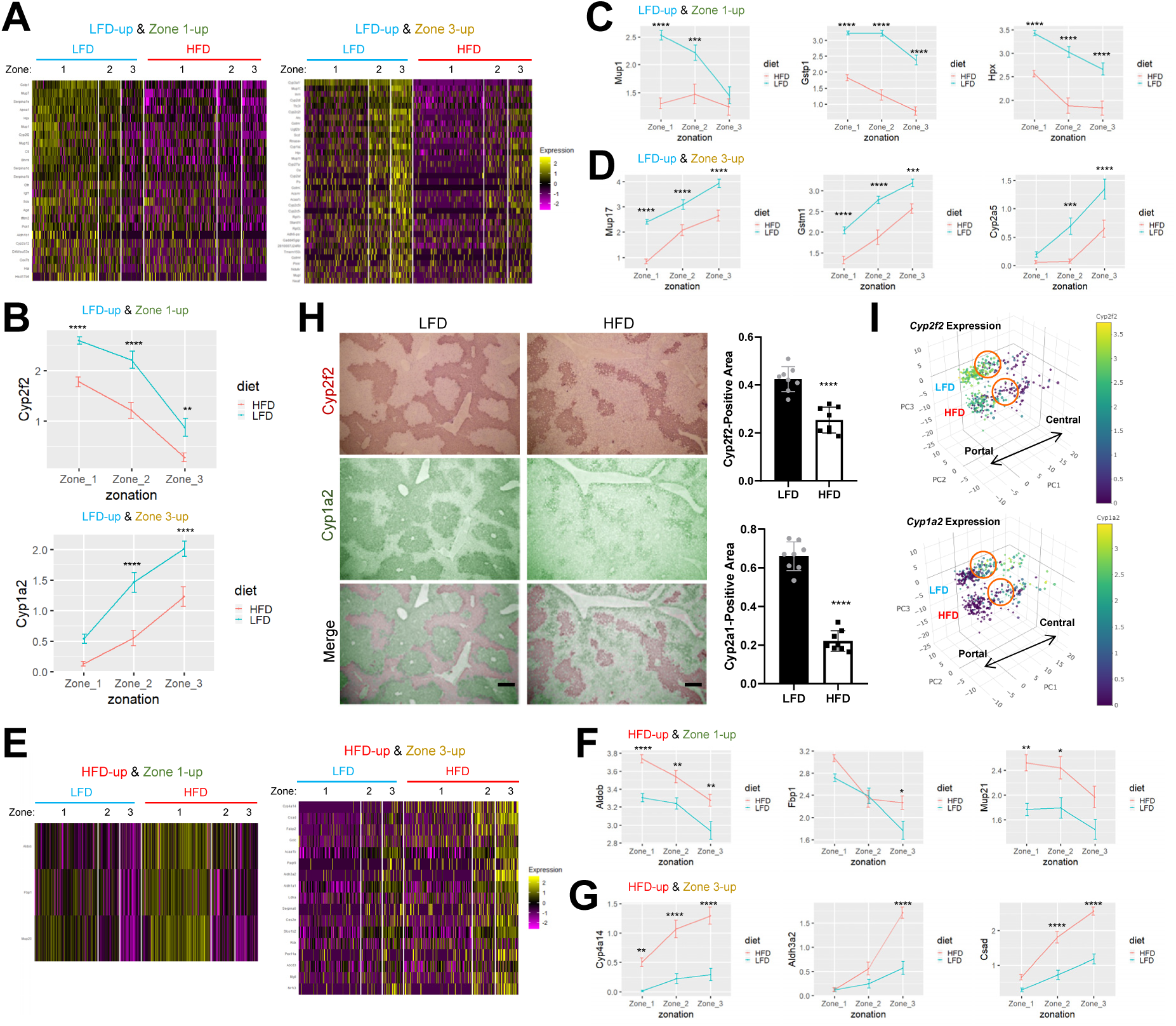
Isolation of genes that are substantially influenced by both diet and zonation. (A and E) Heat map analysis depicting gene expression across single cell population. Cells were clustered into six groups according to diet and zone. HFD-downregulated (A) and upregulated (E) genes that show periportal (zone 1-high; left in each panel) or pericentral (zone 3-high; right in each panel) patterns of expression were analyzed. (B-D, F and G) Analysis of single cell gene expression in hepatocytes of each zone, expressed as mean±SEM (scaled expression values). Data from LFD and HFD livers were analyzed separately. \**P*<0.05, \*\**P*<0.01, \*\*\**P*<0.001, \*\*\*\**P*<0.0001 in Sidak’s multiple comparison test. (H) Cyp2f2 and Cyp1a2 protein expression was visualized through immunohistochemistry from serial sections of LFD and HFD mouse liver (left). Cyp2f2 and Cyp1a2 staining signals were artificially colored with red (first row) and green (second row), respectively, to produce merged images (third row) of the serial liver sections. Cyp2f2- and Cyp1a2-positive areas were quantified (right). Scale bars: 200 µm. (I) 3-dimensional PCA manifold depicting the single cell expression levels of indicated genes. Individual dots represent single cell transcriptome. The size of dots represent the number of RNA features captured in the droplet. LFD and HFD area, as well as the directionality of metabolic zonation (from portal to central), are indicated in each manifold. Orange circles indicate the approximate position of zone 2 hepatocytes.

We assessed whether mRNA expression changes observed from our Drop-seq analysis could lead to alterations of the protein level by examining *Cyp2f2* and *Cyp1a2* genes, which are among the genes that show the strongest zonation patterns in our dataset and previous datasets (1). In our dataset, the expression of these genes in their corresponding metabolic zones was strongly reduced after HFD (Fig. 4B). These observations were reproduced through immunohistochemical staining of Cyp2f2 and Cyp1a2 proteins in liver sections; the areas expressing these two proteins were dramatically shrunken (Fig. 4H). Correspondingly, although the regions expressing Cyp2f2 and Cyp1a2 substantially overlapped in LFD liver, they hardly overlapped in HFD, creating the gap area where none of these proteins were expressed (Fig. 4H). Similar patterns were also observed in Drop-seq data, where many zone 2 hepatocytes reduced expression of both *Cyp2f2* and *Cyp1a2* upon HFD (cells in orange circles of Fig. 4I). These data exemplify the relevance of our Drop-seq dataset for understanding single cell gene expression of hepatocytes in LFD and HFD mouse liver.

### Zonation-independent heterogeneity in single hepatocyte responses to HFD

*Elovl5, Apoa4*, and *Fabp1* are among the genes that show the strongest upregulation of gene expression after HFD (Table S1, third tab). Although the HFD induction of these genes was very robust in both the Drop-seq dataset (Fig. 5A and 5B) and immunohistochemical staining of liver sections (Fig. S5), they did not show strong zone-specific expression patterns (Fig. 5A, 5B and S5). Interestingly, in liver immunohistochemistry, Elovl5, Apoa4, and Fabp1 proteins exhibited variegated expression patterns across the hepatocytes (Fig. S5A), indicating that their induction after HFD is highly heterogeneous between different hepatocytes, independent of metabolic zonation.

**Fig. 5.**
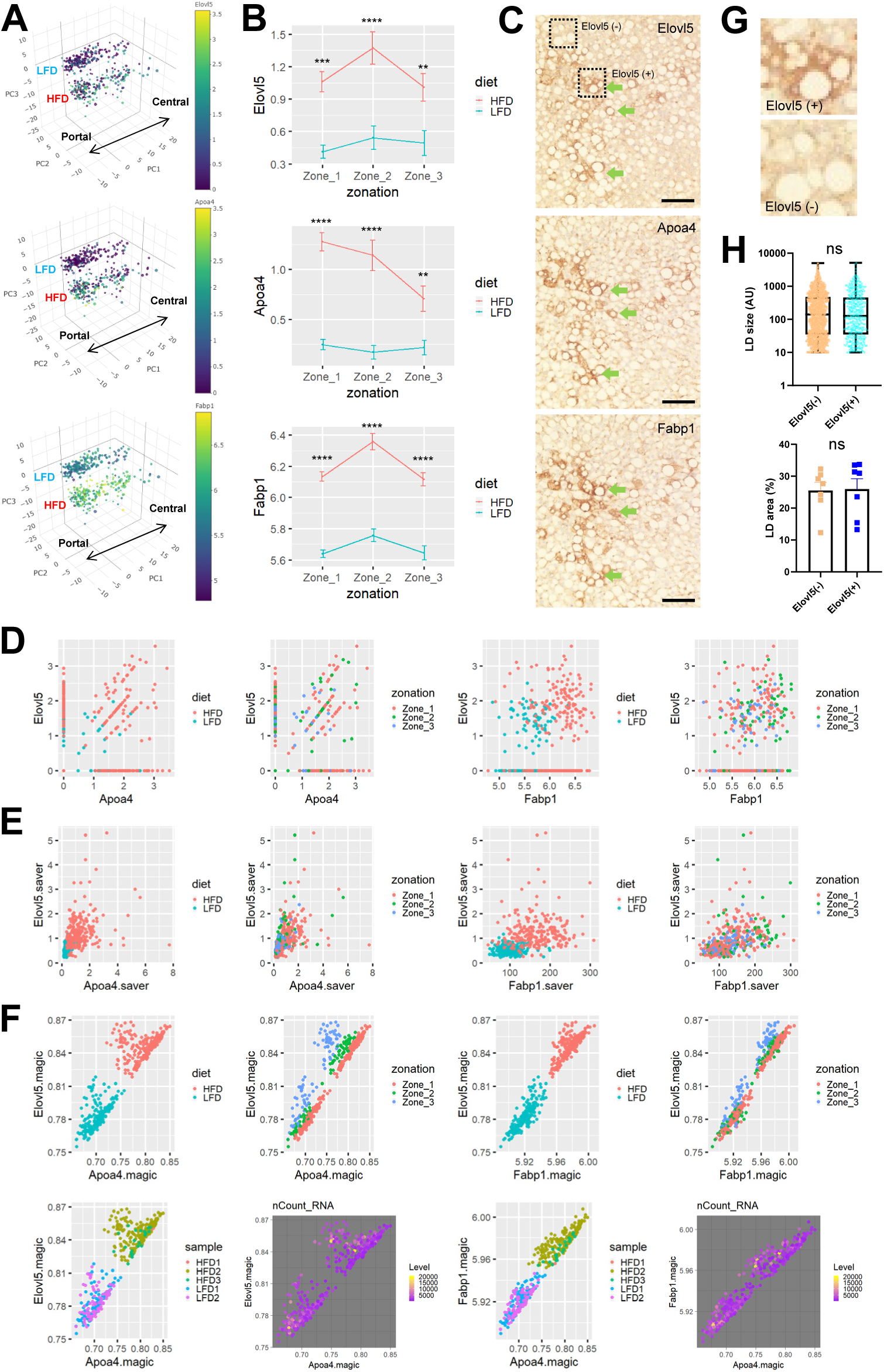
Spatial co-expression pattern of HFD-induced genes regulating lipid metabolism. (A) 3-dimensional PCA manifold depicting the single cell expression levels of indicated genes. Individual dots represent single cell transcriptome. The size of dots represent the number of RNA features captured in the droplet. LFD and HFD area, as well as the directionality of metabolic zonation (from portal to central), are indicated in each manifold. (B) Analysis of single cell gene expression in hepatocytes of each zone, expressed as mean±SEM (scaled expression values). Data from LFD and HFD livers were analyzed separately. \*\**P*<0.01, \*\*\**P*<0.001, \*\*\*\**P*<0.0001 in Sidak’s multiple comparison test. (C, G and H) Elovl5, Apoa4, and Fabp1 protein expression was visualized through immunohistochemistry from serial sections of HFD mouse liver (C). Green arrows indicate areas of positive staining that is congruently observed across all three staining images. Elovl5-high (+) and low (-) areas (dotted boxes) were magnified in (G). Lipid droplet (LD) size (H, upper; n≥479) and area (H, lower; n=7) in Elovl5-high and -low areas were quantified. Data are expressed as a box plot (top; AU, arbitrary unit) or mean±SEM (bottom; % area) with individual data points. Student’s t-tests failed to detect a significant difference between the two groups (ns). Scale bars: 100 µm. (D-F) Correlation between expression levels of *Elovl5, Apoa4*, and *Fabp1* genes from scaled (D), *saver*-imputed (E), and *magic*-imputed (F) Drop-seq dataset. Individual dots represent single cell expression levels, colored by diet, zone, sample information, and level of total RNA counts (nCount_RNA).

Given the spatially-restricted patterns of Elovl5, Apoa4, and Fabp1 protein expression in liver sections (Fig. S5A), we became curious about whether the patterns between these genes are correlated with each other. To assess this, we stained each of these proteins in a serial section of the same histological block. Interestingly, it was found that the regions of high Elovl5, Apoa4 and Fabp1 expression were substantially overlapping with each other, indicating that protein products of these genes are expressed in a positive correlation with each other (Fig. 5C).

We then examined whether the positive correlation between *Elovl5, Apoa4*, and *Fabp1* expression could be observed from the Drop-seq dataset. Query of the most significantly correlated gene for *Elovl5* expression resulted in *Apoa4, Cyp4a14*, and *Fabp1* as the top 3 genes, among which both *Apoa4* and *Fabp1* are included. Correlation scatterplot between *Elovl5* and these two genes showed the trend of positive correlation in scaled data (Fig. 5D; *r* = 0.22 and 0.21, respectively; *P* < 1.5e-6); however, due to the sparsity of the specific mRNA observation and subsequent technical noise, the observed correlation may not be as strong as the true correlation. After applying two independent imputation methods correcting for the technical noise effect, *saver* (17) (Fig. 5E) and *magic* (47) (Fig. 5F), we were able to detect more robust correlation between gene expression profiles of *Elovl5* and *Apoa4* (*r* = 0.90 (magic) and 0.50 (saver)), and between *Elovl5* and *Fabp1* (*r* = 0.97 (magic) and 0.41 (saver)) (Fig. 5E and 5F). Importantly, these patterns of correlation were independent of the zonation (Fig. 5D-5F, zonation panels), batches (Fig. 5F, sample panels), or mRNA reads (Fig. 5F, nCount_RNA/level panels). Therefore, these results suggest the presence of zonation-independent heterogeneity in hepatocyte responses to HFD.

### Elovl5-high and -low hepatocytes accumulate similar levels of fat droplets

In HFD mice, hepatocytes expressing high levels of Elovl5, Apoa4, and Fabp1 were not morphologically different from other hepatocytes in terms of lipid droplet accumulation (Fig. 5C and 5G). Quantification of the lipid droplet size did not reveal any obvious differences in lipid droplet size (Fig. 5H, upper) or area (Fig. 5H, lower) between Elovl5-high and -low hepatocyte populations. Therefore, the levels of HFD-induced Elovl5, Apoa4, and Fabp1 does not seem to substantially alter the steady-state level of fat accumulation in the hepatocytes.

Considering that *Fabp1, Elovl5* and *Apoa4* are all involved in fatty acid metabolism, it could be inferred that hepatocytes expressing high levels of these genes might be more active in lipid processes. Since the histological analysis indicates that the expression of these genes does not substantially alter the intracellular amounts of fat droplets (Fig. 5G and 5H), the biological relevance of this heterogeneous gene expression pattern is unclear in the context of HFD feeding. It is possible that active processes in lipid metabolism, mediated by these genes, alter the flux of lipid metabolites without affecting the steady-state fat levels. It is also possible that heterogeneity in expression of these genes is temporarily generated; therefore, over time, other hepatocytes might also express high levels of these proteins, producing similar metabolic profiles.

### HFD induces stronger fat accumulation in zone 3 hepatocytes

It was well documented that obesity and hepatosteatosis disparately affect individual hepatocytes across their histological zonation. Some hepatocytes, especially the ones in zone 3, which are deprived of nutrients and oxygen, are more prone to accumulate lipid droplets while the ones in zone 1, a nutrient- and oxygen-rich environment, are less susceptible to steatotic progression (14, 15). Consistent with these former studies, we observed from the histology results that Cyp1a2-positive zone 3 hepatocytes contain more and bigger lipid droplets when compared to Cyp2f2-positive zone 1 hepatocytes (Fig. 6A and 6B). The observation of zone 3-specific fat accumulation was reproduced when we directly stained lipid droplets in freshly frozen tissue sections from LFD and HFD livers (Fig. 6C), again supporting that HFD-induced fat accumulation is more pronounced in zone 3.

**Fig. 6.**
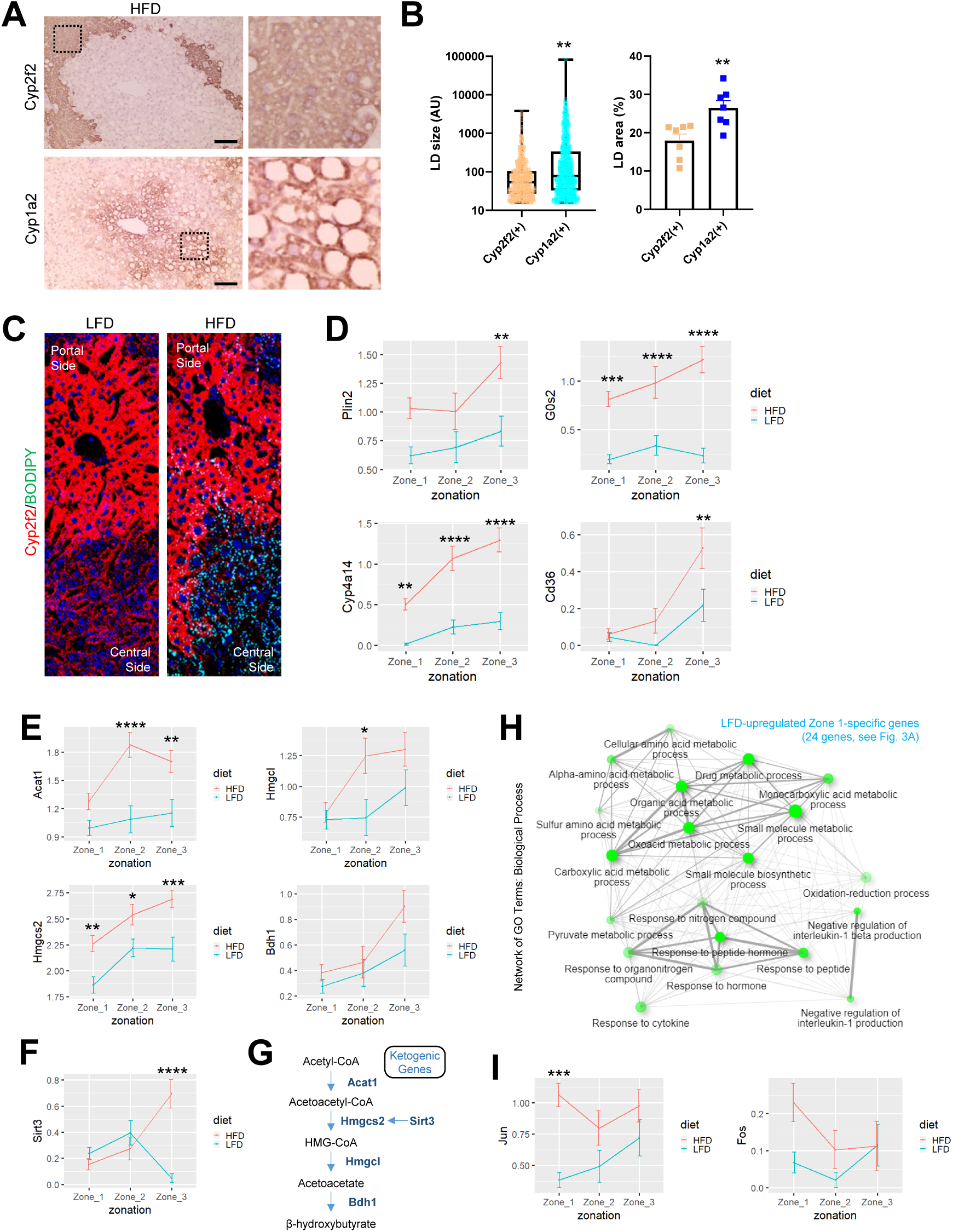
HFD induces zone 3 hepatocytes to express genes promoting lipid droplet accumulation. (A and B) Serial sections of HFD mouse liver were stained with zone 1 marker Cyp2f2 (A, upper) and zone 2 marker Cyp1a2 (A, lower). Boxed areas were magnified in the right (A). LD size (B, left; n≥535) and area (B, right; n=7) in each compartment was quantified. Data are expressed as box plot (left; AU, arbitrary unit) or mean±SEM (right; % area) with individual data points. Student’s t-tests were used to examine significant difference between the two groups (\*\**P*<0.01). Scale bars: 100 µm. (C) Fresh frozen sections from LFD and HFD mouse liver were immunostained to visualize Cyp2f2 (red), lipid droplets (green, stained by BODIPY 493/503) and DNA (blue, by DAPI). (D-F and I) Analysis of single cell gene expression in hepatocytes of each zone, expressed as mean±SEM (scaled expression values). Data from LFD and HFD livers were analyzed separately. \**P*<0.05, \*\**P*<0.01, \*\*\**P*<0.001, \*\*\*\**P*<0.0001 in Sidak’s multiple comparison test. (G) Ketogenic genes induced by HFD in zone 3 hepatocytes are presented in a pathway diagram. (H) Network analysis of gene ontology-biological pathway (GO-BP) enrichment terms in the HFD-downregulated zone 1-specific gene list, using ShinyGO (11). Pathways whose enrichment is significant (FDR < 0.05) were presented as nodes. Two nodes are connected if they share 20% or more genes. Darker nodes are more significantly enriched gene sets. Bigger nodes represent larger gene sets. Thicker edges represent more overlapped genes.

### Zone 3 hepatocytes robustly express genes mediating fat accumulation during HFD

We then tried to identify the features of single hepatocyte transcriptome that may explain the preferential accumulation of lipid droplets in zone 3 hepatocytes. For this, we surveyed the function of all genes that show either HFD- or zone 3-specific expression patterns (Table S1, third and fifth tabs) through literature search. We found that there are at least four genes, *Plin2, G0s2, Cyp4a14*, and *Cd36*, which are known to play a mechanistic role in fat accumulation (23, 26, 31, 50, 54-56), and at the same time, strongly induced by HFD in zone 3 hepatocytes (Fig. 6D).

Plin2 is a protein directly associated with hepatic lipid droplets (26). Plin2 surrounds the lipid droplet and assists the storage of neutral lipids within the lipid droplets. Consistent with increased lipid droplet accumulation in zone 3 hepatocytes, *Plin2* expression is more strongly upregulated in zone 3 during HFD (Fig. 6D). The *Plin2* induction could be critical for zone 3-specific accumulation of lipid droplets because hepatic deletion of *Plin2* is known to attenuate hepatic fat accumulation (26, 31).

*G0s2*, whose product is a well-established inhibitor of lipase activity in hepatocytes (54), was also strongly upregulated upon HFD, specifically in zone 3 hepatocytes (Fig. 6D). Considering that G0s2 is important for the accumulation of triglycerides in hepatocytes by inhibiting lipase activities, it is likely that the pericentral expression of *G0s2* is responsible for lipid droplet accumulation in zone 3. Indeed, in a recent study, *G0s2* knockout mice and liver-specific knockdown mice did not show hepatosteatosis upon HFD, while G0s2 overexpression sufficed to induce hepatosteatosis (56).

*Cyp4a14* is another gene that is induced upon HFD and critical for generating HFD-induced hepatosteatosis (55). HFD-induced *Cyp4a14* expression is also much more pronounced in zone 3, compared to the other zones (Fig. 6D).

*Cyp4a14* was suggested to promote hepatosteatosis, partly thorough inducing *Cd36/FAT*, whose products play a role in importing fatty acids into hepatocytes (55). *Cd36/FAT* was also highly induced in zone 3 hepatocytes of HFD-fed mouse liver (Fig. 6D). Notably, prior studies showed that hepatic *Cd36* overexpression was sufficient to provoke hepatosteatosis even without HFD challenges (23), while liver-specific *Cd36* disruption was sufficient to attenuate fatty liver in HFD mice (50).

Collectively, these observations, combined with former genetic studies performed on these genes (23, 26, 31, 50, 54-56), suggest that zone 3-specific upregulation of *Plin2, G0s2, Cyp4a14*, and *Cd36* plays an important role for producing zone 3-specific steatosis phenotype in response to HFD challenges.

### Zone 3 hepatocytes upregulate genes mediating ketogenic pathway

In addition to the genes responsible for producing lipid droplet accumulation, we also observed that HFD-induced expression of ketogenic genes, such as *Acat1, Hmgcs2, Hmgcl* and *Bdh1*, were relatively higher in zone 3 hepatocytes (Fig. 6E). *Sirt3*, whose product deacetylates and activates Hmgcs2, was also more strongly expressed in zone 3 hepatocytes of HFD liver (Fig. 6F). These results suggest that the HFD-induced ketogenesis pathway (Fig. 6G) is preferentially activated in zone 3 hepatocytes of mouse liver. Activation of ketogenesis in zone 3 hepatocytes might be critical for distributing the energy to peripheral tissues and generating metabolic adaptation to HFD-induced hypernutrition (36).

### Zone 1 hepatocytes also transcriptionally respond to HFD

In contrast to zone 3 hepatocytes, zone 1 hepatocytes strongly downregulated many zone 1-specific transcripts that mediate various metabolic processes, including drug and amino acid catabolism and redox metabolism (Fig. 6H). Reduction of these functions may be critical for HFD adaptation by accommodating an increased need for lipid metabolism. Although many of zone 1-specific genes were downregulated (Fig. 4A), *Aldob* and *Fbp1*, two genes that are involved in gluconeogenesis, were strongly upregulated in zone 1 hepatocytes after HFD (Fig. 4E and 4F). This is consistent with the previous findings indicating that gluconeogenesis activity is the most active in zone 1 hepatocytes (15). This zone 1-specific regulation of *Aldob* and *Fbp1* might be contributing to decreased glucose tolerance during HFD-induced obesity (51). In addition, stress-induced AP1 transcription factors, *Jun* and *Fos*, were also specifically upregulated in zone 1 hepatocytes upon HFD stimulation (Fig. 6I). These results indicate that, although zone 1 hepatocytes are relatively resistant to fat droplet accumulation, they also respond transcriptionally to HFD challenges and contribute to physiological HFD responses in a substantial way.

### PPAR pathway is implicated in HFD modulation of single hepatocyte transcriptome

It is interesting to note that many HFD-induced genes reviewed above are targets of the PPAR-family transcription factors; genes with a variegated co-expression pattern (*Elovl5, Apoa4*, and *Fabp1*), as well as genes that show zone 3-specific pattern and mediate fat droplet accumulation (*G0s2, Plin2, Cyp4a14*, and *Cd36*) and ketogenesis upregulation (*Acat1, Hmgcs2, Hmgcl* and *Bdh1*), are all targets or PPARα (21, 29, 49) (Fig. 7). As observed above (Fig. 3E-G), the PPAR pathway is the only transcription factor-targeted group that is enriched in both HFD-upregulated and HFD-downregulated gene lists. Importantly, PPAR is known to be activated upon stimulation with fatty acids, as they are direct ligands for transcriptional activation of PPAR (21, 29). Former bulk analysis of fatty liver transcriptome also revealed that various targets of PPAR, as represented in our dataset, are strongly upregulated upon HFD challenges (22, 33, 37, 39). Notably, many of these genes did not show such diet-dependent modulations in *Ppara*-deleted knockout mutant strains (33, 49). Therefore, many transcriptome features observed from our dataset could be at least partly mediated by PPAR activation by excessive fatty acids from dietary sources.

**Fig. 7.**
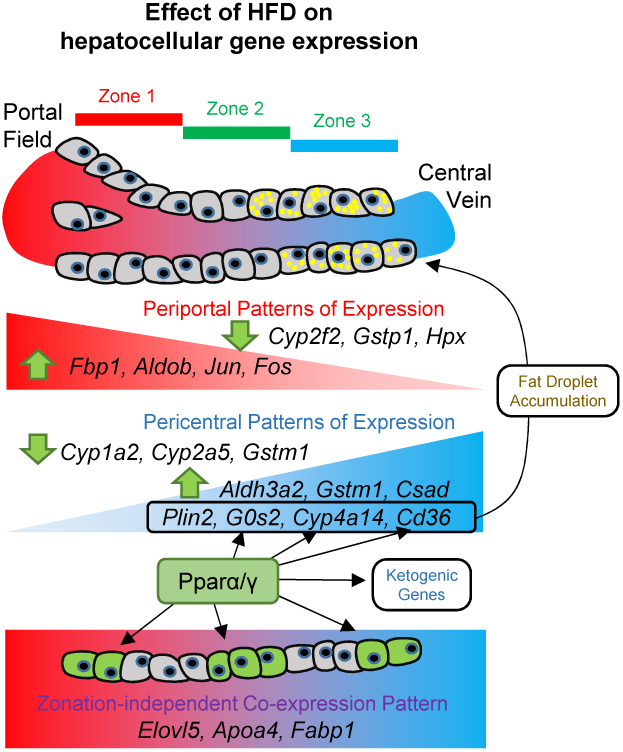
Holistic understanding of heterogeneous hepatocyte responses to HFD. The schematic diagram depicts the heterogeneous effect of HFD on single hepatocellular gene expression. Our dataset indicates that the entire hepatocytes population undergoes substantial transcriptome changes upon HFD, and that the patterns of alteration were highly heterogeneous across the hepatocyte population with zonation-dependent and -independent effects.

### Limitations of the current study

Our Drop-seq dataset contains a large number of droplets containing ambient RNA (Fig. S1, see Materials and Methods for details), indicating that a considerable number of single hepatocytes were damaged during isolation and microfluidics. It is possible that the single hepatocyte data presented in the current study is biased towards the hepatocyte population that is resistant to physical damage. In addition, although our method of partitioning single hepatocyte transcriptome profiles into three zones is robust and consistent with previous studies, it is possible that this is an oversimplification of the complex histological architecture of the liver. These issues could be potentially addressed in future studies by utilizing microfluidics-free methods for sorting single cells (1, 13) or spatial profiling of liver transcriptome through tissue sections (35).

## Summary

Recent single cell transcriptome studies revealed that hepatocyte gene expression and function are highly heterogeneous across their metabolic zonation, revealing global division of metabolic labor of the liver (1, 28). Building on these previous findings, our study provides the first snapshot of how single hepatocyte transcriptome landscape is altered in response to HFD and subsequent development of NAFLD. Through this dataset, we were able to find that HFD makes an impact on the transcriptome of the entire hepatocyte population. We also found that HFD responses of hepatocytes can be heterogeneous with zonation-dependent and - independent effects. Our observations detailed above systematically characterize HFD-induced changes in hepatocellular transcriptome and their relationship to NAFLD pathogenesis. Furthermore, we made our dataset available in an interactive web tool (https://lee.lab.medicine.umich.edu/hfd), where individual investigators can reproduce our analyses and test their hypothesis using our publically available dataset. We believe that this resource will be greatly useful for future NAFLD studies.

## Acknowledgment

We thank Drs. Jiandie Lin, Yatrik Shah, and Lei Yin labs for their help in hepatocyte isolation and scRNA-seq optimization. We thank Dr. Myungjin Kim for advice, Dr. Steve Norris for editing, Bondong Gu and Boyoung Kim for assistance, and Santa Cruz Biotechnology for antibodies.

## Grants

This work was supported by the NIH (R01DK102850 and R01DK114131 to J.H.L., U01HL137182 to H.M.K., T32AG000114 to C.S.C. and P30AG024824, P30DK034933, P30DK089503, and P30CA046592), the Chan Zuckerberg Initiative (to H.M.K.), the MCubed initiative (to J.H.L. and H.M.K.), and the AASLD Pilot Research Award (to J.H.L. and H.M.K.).

## Disclosures

No conflicts of interest, financial or otherwise, are declared by the authors.

## Author Contributions

S.R.P. performed Drop-seq experiments. C.S.C. performed histology experiments. J.X. helped with computational analysis. H.M.K. and J.H.L. conceived and directed the project. S.R.P., H.M.K., and J.H.L. designed experiments, analyzed data, and wrote the manuscript. All authors approved the final version.

## Notes

### Competing Interest Statement

The authors have declared no competing interest.

### Summary of Updates

Results and Discussion were combined. Text and Figures were reorganized and updated.

