## Supplementary figures and images for "Holistic Characterization of Single Hepatocyte Transcriptome Responses to High Fat Diet"

### Supplemental Fig S1-S5

Park et al. Figure S1

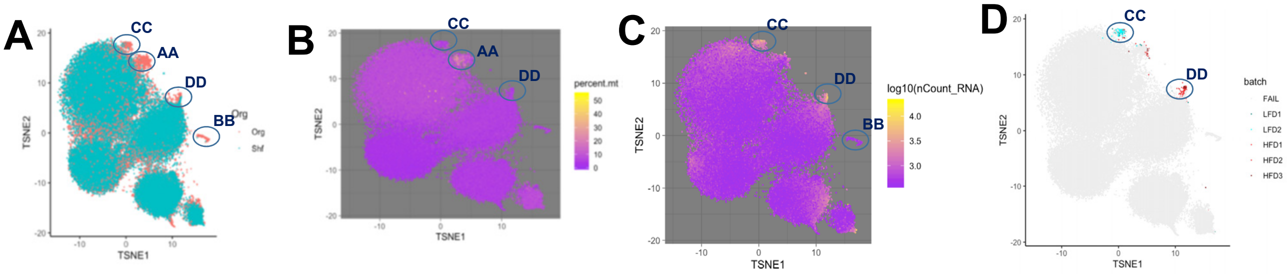

Park et al. Figure S2

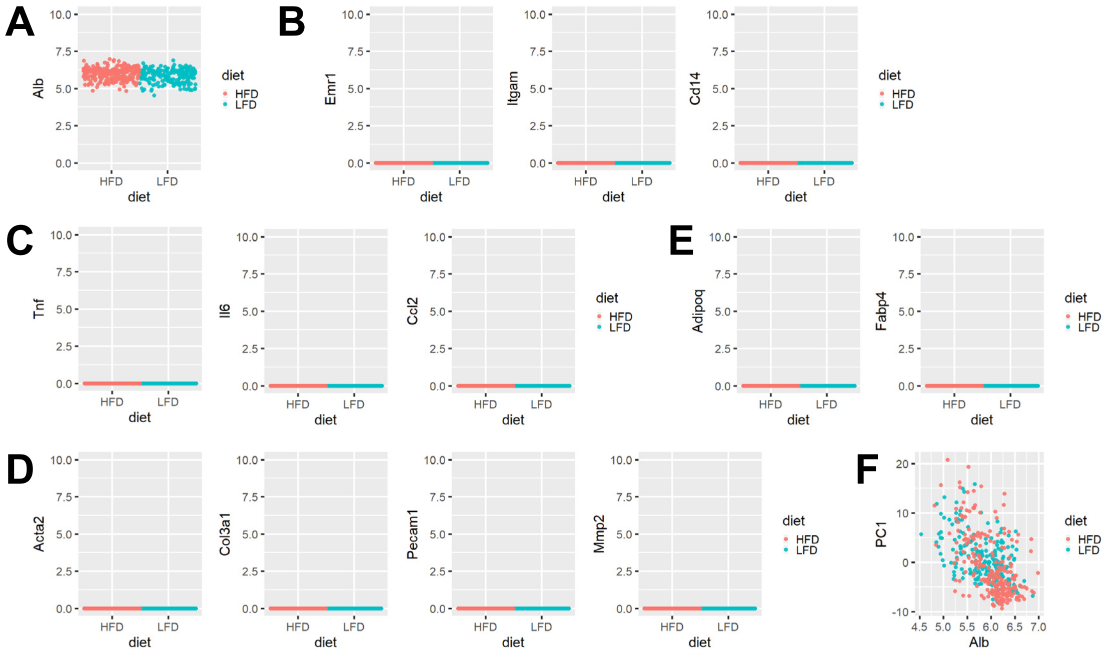

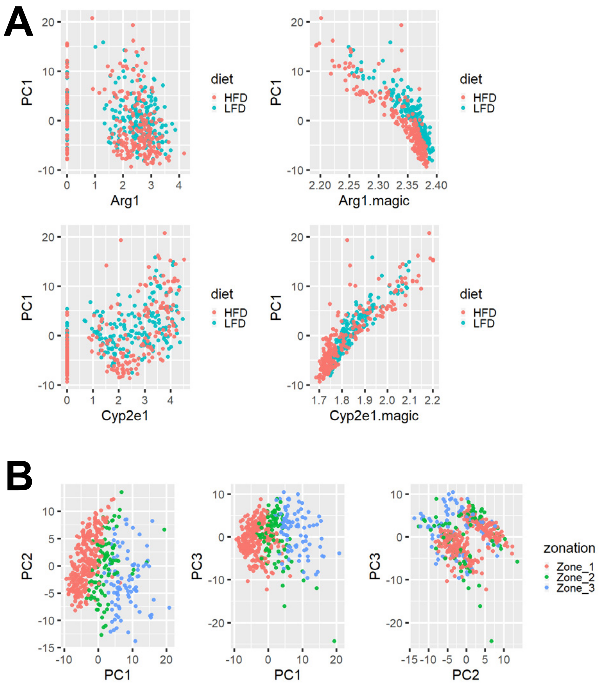

Park et al. Figure S4

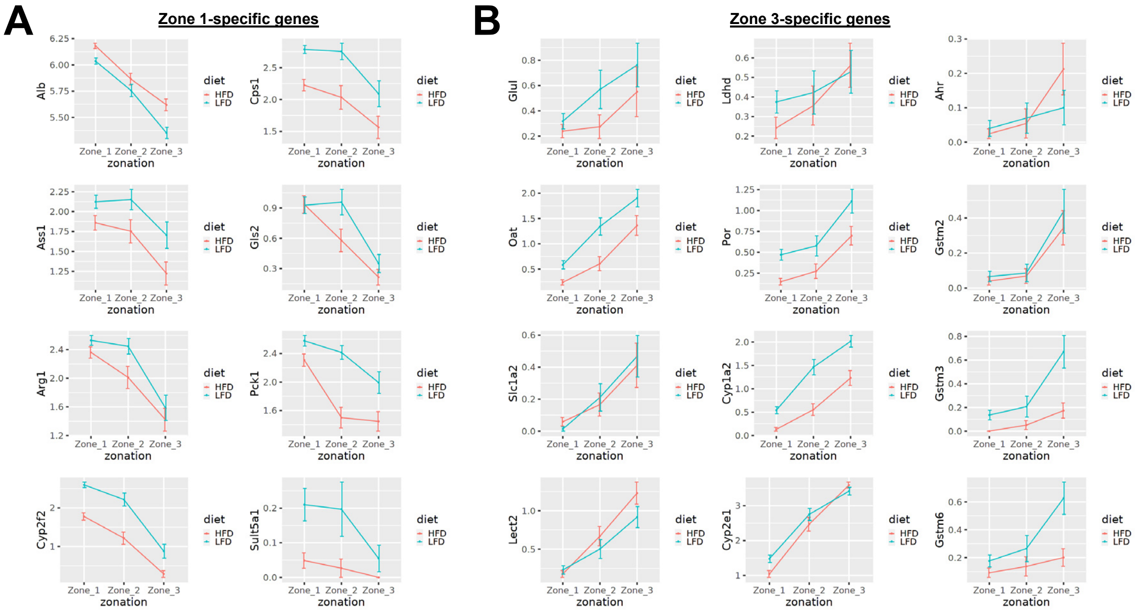

Park et al. Figure S5

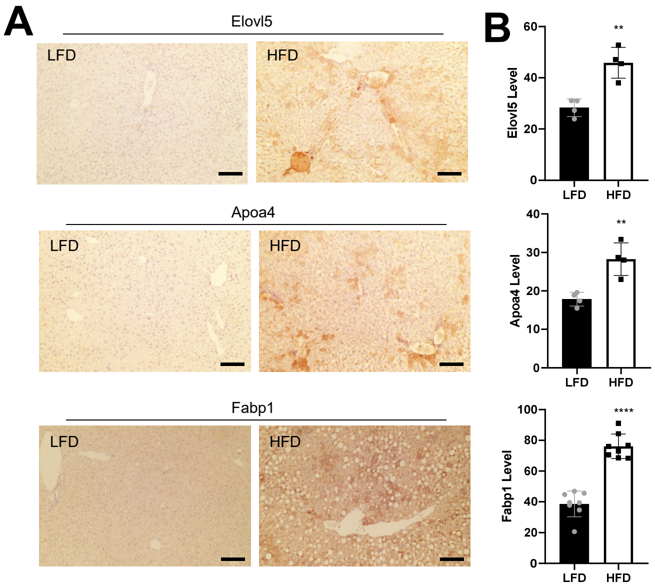
