## Supplemental Fig/Table Legend for "Holistic Characterization of Single Hepatocyte Transcriptome Responses to High Fat Diet"

**Supplemental Figure Legends**

**Fig. S1-S5**

**Supplemental Table Legend**

**Table S1**

**Supplemental Figure Legends**

**Fig. S1.** Isolation of single hepatocyte transcriptome data from the original Drop-seq dataset.
(A) t-SNE manifold of original Drop-seq dataset (Org, red) and a shuffled dataset (Shf, blue) that was produced by random shuffling of gene expression data in the original dataset. Individual dots represent gene expression data in each droplet from Drop-seq. Four major areas (AA-DD) were identified as potential droplet clusters that represent real single cell hepatocyte transcriptome, because they do not overlap with clusters that contain randomly shuffled droplets.
(B) t-SNE manifold depicting single droplet expression of mitochondrial genes. Cluster AA contains a very high amount of mitochondrial transcripts, which is indicative of droplets containing dead cells or cell debris.
(C) t-SNE manifold depicting how many different RNAs were captured from the droplets. Cluster BB contained a low number of RNAs indicating that these droplets do not contain real cells.
(D) Through a series of multi-dimensional clustering, CC and DD clusters were isolated from the total Drop-seq dataset, which represent single hepatocytes form LFD and HFD mouse liver, respectively. All experimental batches were represented in the CC and DD clusters.

**Fig. S2.** Hepatocyte specificity of the current Drop-seq dataset.
(A-D) Single cell expression levels of each indicated gene (scaled expression values), analyzed in LFD and HFD livers, represented in each dot in the graph. (A) Hepatocyte marker *Alb*, (B) macrophage markers, (C) inflammatory markers, (D) hepatic stellate cell markers, and (E) adipocyte markers.
(F) Negative correlation between *Alb* expression and the PC1 level. Individual dots, colored by diet, represent single cell transcriptome and levels on each axis.

**Fig. S3.** Identification of zonation patterns in single cell transcriptome of LFD and HFD livers.
(A) Correlation between the PC1 level and indicated gene expression levels. Scaled (left) and *magic*-imputed (right) levels of *Arg1* (upper) and *Cyp2e1* (lower) expression were used for analyses. PC1 is negatively correlated with *Arg1* and positively correlated with *Cyp2e1*, suggesting that it represents portal-to-central zonation patterns.
(B) 2-dimensional PCA manifolds depicting zone identity, determined by imputed levels of *Arg1* and *Cyp2e1* expression.

**Fig. S4.** Validation of zone layering through established zonation marker genes.
(A and B) Analysis of single cell gene expression in hepatocytes of each zone, layered according to our method, expressed as mean±SEM (scaled expression values). Established markers for zone 1 (A) and zone 3 (B) hepatocytes were analyzed. Data from LFD and HFD livers were analyzed separately.

**Fig. S5.** Elovl5, Apoa4 and Fabp1 show variegated patterns of expression across liver sections.
(A and B) Indicated protein expression was visualized through immunohistochemistry from LFD and HFD mouse livers (A). Immunostaining intensities were quantified (B; n≥4). Data are expressed as mean±SEM with individual data points (AU, arbitrary intensity unit). Student’s t-tests were used to examine significant difference between the two groups (***P*<0.01, **** *P*<0.0001). Scale bars: 100 µm.

**Supplemental Table Legend**

**Table S1.** List of genes that show diet- and zone-specific expression patterns in single hepatocyte transcriptome analysis.
(Tab 1: SUMMARY) List of LFD-upregulated (LFD-up), HFD-upregulated (HFD-up), zone 1-upregulated (Portal-up), zone 3-upregulated (Central-up), LFD- and zone 1-upregulated (LF-Portal), LFD- and zone 3-upregulated (LF-Central), HFD- and zone 1-upregulated (HF-Portal), and HFD- and zone 3-upregulated (HF-Central) genes.
(Tab 2: LFD-UP) List of LFD-upregulated genes, their significance of LFD-specific expression (p_val) and average fold change above the background level (avg_logFC).
(Tab 3: HFD-UP) List of HFD-upregulated genes, their significance of HFD-specific expression (p_val) and average fold change above the background level (avg_logFC).
(Tab 4: Portal-UP) List of zone 1-upregulated genes, their significance of zone 1-specific expression (p_val) and average fold change above the background level (avg_logFC).
(Tab 5: Central-UP) List of zone 3-upregulated genes, their significance of zone 3-specific expression (p_val) and average fold change above the background level (avg_logFC).
